# Loss of a novel striated muscle-enriched mitochondrial protein Coq10a enhances postnatal cardiac hypertrophic growth

**DOI:** 10.1101/755793

**Authors:** Kentaro Hirose, Steven Chang, Hongyao Yu, Jiajia Wang, Emanuele Barca, Xiaoxin Chen, Sheamin Khyeam, Kazuki Tajima, Takeshi Yoneshiro, Shingo Kajimura, Catarina M. Quinzii, Guang Hu, Guo N. Huang

**Affiliations:** Cardiovascular Research Institute & Department of Physiology, University of California, San Francisco, San Francisco, CA, 94158, USA; Eli and Edythe Broad Center for Regeneration Medicine and Stem Cell Research, University of California, San Francisco, San Francisco, CA, 94158, USA; Epigenetics and Stem Cell Biology Laboratory, National Institute of Environmental Health Sciences, RTP, NC 27709, USA; Department of Neurology, Columbia University Medical Center, New York, NY, United States; UCSF Diabetes Center, San Francisco, CA, USA; Department of Cell and Tissue Biology, University of California, San Francisco, CA, USA

## Abstract

Postnatal mammalian cardiomyocytes undergo a major transition from hyperplasia (increases in cell numbers) to hypertrophy (expansion in cell size). This process is accompanied by rapid mitochondrial biogenesis and metabolic switches to meet the demand of increased cardiac output. Although most mitochondrial components express ubiquitously, recent transcriptomic and proteomic analyses have discovered numerous tissue-specific mitochondrial proteins whose physiological functions are largely unknown. Here we report that a highly evolutionarily conserved mitochondrial protein Coq10a is predominantly expressed in mammalian cardiac and skeletal muscles, and is highly up-regulated around birth in a thyroid hormone-dependent manner. Deletion of *Coq10a* by CRISPR/Cas9 leads to enhanced cardiac growth after birth. Surprisingly, adult *Coq10a* mutant mice maintain the hypertrophic heart phenotype with increased levels of coenzyme Q (CoQ) per cardiomyocyte, preserved cardiac contractile function and mitochondrial respiration, which contrasts with reported mice and humans with mutations in other Coq family genes. Further RNA-seq analysis and mitochondrial characterization suggest an increase of mitochondrial biogenesis in the *Coq10a* mutant heart as a possible consequence of Peroxisome proliferator-activated receptor Gamma Coactivator 1-alpha (PGC1α) activation, consistent with a recent intriguing report that CoQ may function as a natural ligand and partial agonist of Peroxisome Proliferator-Activated Receptor (PPAR) α/γ. Taken together, our study reveals a previously unknown function of a novel striated muscle-enriched mitochondrial protein Coq10a in regulating postnatal heart growth.

## Introduction

As highly evolutionarily conserved electron carrier in all eukaryotes, the lipid coenzyme Q (CoQ, ubiquinone), plays a pivotal role in the electron transport chain by carrying electrons from mitochondrial respiratory chain complexes I and II to complex III (*1–3*). The synthesis and transport of CoQ require orchestrated actions of a complex of enzymes and ancillary proteins (*2, 3*). In yeast, the genes encoding coenzyme Q-synthase complex proteins (*COQ1*-*11*) have been identified from mutants with reduced CoQ levels and defective mitochondrial respiration, and some of these mutant phenotypes can be rescued by supplementation of CoQ (*1*). Among these Coq proteins, Coq1p-Coq9p proteins are known to be involved in CoQ synthesis (*1*).

Yeast *COQ10* gene belongs to the Coq family and as in other Coq family mutants, *coq10Δ* mutants have deficient mitochondrial respiration (*4*). However, yeast *coq10Δ* mutants show nearly normal levels of CoQ at stationary phase unlike other coq family mutants that lack CoQ (*4, 5*). Coq10p contains a domain termed START that is implicated in lipid and sterol binding and transport (*4*). Indeed, Coq10p derived from yeasts biochemically binds to CoQ (*4, 6*). Thus, Coq10p is suggested to function as a carrier of CoQ to its proper location in the mitochondria. Mammalian Coq10a is known as a homolog of yeast Coq10p because human COQ10A contains the key CoQ-binding amino acids and can rescue mitochondrial respiration defects in yeast *coq10Δ*(*4, 7, 8*). However, its physiological function in mammalian tissues has never been investigated previously.

CoQ was initially isolated from mitochondria in beef heart (*9*), and heart is the tissue that contains one of the highest levels of CoQ (*10*). Patients or rodent models with mutations in various mitochondrial genes including Coq family genes often develop cardiomyopathy (*11*). It has been reported that human patients carrying mutations in CoQ synthesis genes (*PDSS1*, *PDSS2, COQ2*, *COQ4*, *COQ7*, *COQ8B*, *COQ9*) lead to cardiac defects including hypertrophic cardiomyopathy and dilated cardiomyopathy (*2, 12*). Analyses of 5,000 knockout mouse lines by the International Mouse Phenotype Consortium revealed that knockout of the mouse homolog of *COQ2*, *COQ4*, *COQ6*, *COQ8B*, *PDSS1* or *PDSS2* in mice are homozygous-lethal (*13*), highlighting the essential role of CoQ synthesis in development. A mouse model *Coq9^R239X^* harboring a premature stop codon found in a human patient presents widespread tissue deficiency of CoQ and develops signs of cardiac fibrosis (*14*). So far, mutations in the *COQ10A* gene have never been reported in humans. Here we show that rodent *Coq10a* (a homolog of human *COQ10A*) is predominantly expresses in the heart muscle mitochondria. The mRNA level of *Coq10a* is positively regulated by thyroid hormone signaling in postnatal mouse heart and isolated cardiomyocytes. To understand the physiological function of *Coq10a*, we generated *Coq10a* knockout mice using the CRISPR/Cas9-based genome editing technique. *Coq10a* mutant mice showed cardiac hypertrophic phenotype without any features of pathological hypertrophy such as fetal gene expression, cardiac dysfunction and fibrosis. Furthermore, *Coq10a* knockout hearts have intact mitochondrial respiration and contain higher levels of CoQ, indicating that although mouse Coq10a can rescue the lethal phenotype in yeast *coq10* mutants, it has unique functions in the mammalian heart that are distinct from yeast Coq10p.

## Results

### Coq10a is a novel mitochondrial protein predominantly expressed in postnatal striated muscles

We analyzed a rat RNA-seq transcriptomic BodyMap across 11 organs (*15*) and revealed that *Coq10a* is predominantly expressed in the skeletal muscle and heart at 21 weeks of age (Figure 1A) while its paralog *Coq10b* is highly expressed in the adrenal gland but present in most other organs at a low level (Figure S1A). In addition, a recent detailed analysis of gene expression across mouse organ development showed that cardiac expression of *Coq10a* begins around E15.5, rapidly increases postnatally and keeps increasing by postnatal day 63 (P63) (Figure 1B); *Coq10b* expression follows a similar temporal pattern in the heart (Figure S1B) (*16*). In the adult mouse heart, *Coq10a* expression is 4-5 folds higher than *Coq10b* (Figure 1B and S1B). Consistently, we found that Coq10a protein levels increase in the mouse heart during postnatal development (Figure 1C). Because yeast Coq10p protein localizes to the inner mitochondrial membrane (*4, 6*), we investigated whether mouse Coq10a is also a mitochondrial protein. Subcellular fractionation of the adult mouse ventricular tissue indeed showed that Coq10a is highly enriched in the mitochondrial fraction rather than the cytosolic fraction (Figure 1C). Altogether, these results support Coq10a as a striated-muscle enriched mitochondrial protein.

**Figure 1.**
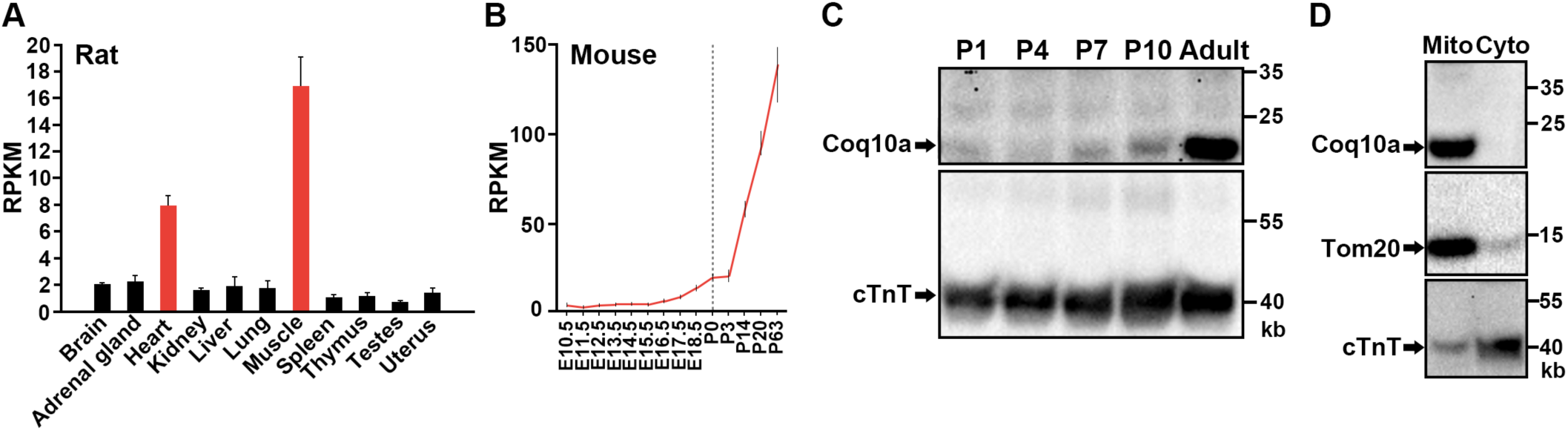
Tissue-specific and temporal expression of *Coq10a* gene in rodents. (**A**) *Coq10a* expression in various adult (21-week old) rat organs based on the transcriptomic BodyMap (*14*). (**B**) Temporal *Coq10a* expression from embryonic day (E) 10.5 to postnatal day (P) 63 in the mouse heart, based on RNA-seq gene expression profile (15). (**C**) Western blotting analysis of Coq10a protein in P1, P4, P7, P10 and adult mouse hearts. A cardiomyocyte cytoskeleton protein cardiac troponin T (cTnT) is used as a loading control. (**D**) Western blotting analysis of Coq10a protein in the mitochondrial (Mito) and cytosolic (Cyto) fractions of adult mouse heart lysates. Tom20 is a marker for mitochondria and cTnT is used as a cytosolic marker.

### Thyroid hormone signaling activates *Coq10a* gene expression

Because the temporal dynamics of *Coq10a* expression matches that of circulating thyroid hormones (*17*) and expression of downstream target genes such as *Myh6* (*18*), we hypothesized that *Coq10a* gene expression is controlled by thyroid hormone signaling in the heart. Thyroid hormone receptors are transcription factors that bind DNA directly and regulate the expression of target genes.

We first tested whether increase of thyroid hormones is sufficient to activate *Coq10a* expression in primary mouse neonatal cardiomyocytes. Supplementation of culture media with triiodothyronine (T3) enhanced the gene expression of *Coq10a* by ∼2.5 folds in primary cardiomyocytes (Figure 2A). Thus, these results suggest that activation of thyroid hormone signaling induces *Coq10a* gene expression.

**Figure 2.**
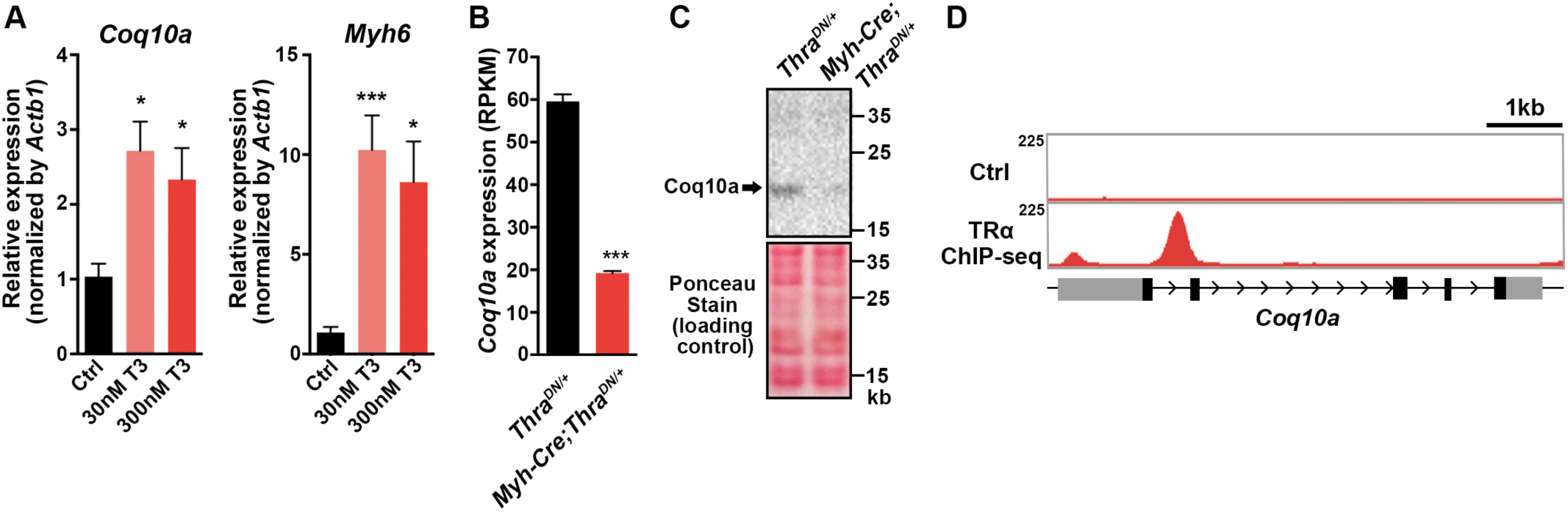
Thyroid hormone signaling directly controls *Coq10a* expression. (**A**) Relative gene expression of *Coq10a* and *Myh6* in triiodothyronine (T3)-treated neonatal mouse cardiomyocytes in culture. (**B**) Gene expression of Coq10a on cardiac-specific thyroid hormone receptor α dominant negative mutant (*Myh6-Cre; Thra^DN/+^*) and littermate control (*Thra^DN/+^*) hearts at P14, base on the RNA-seq analysis. (**C**) Western blotting analysis of Coq10a protein in thyroid hormone signaling deficient (*Myh6-Cre; Thra^DN/+^*) and control (*Thra^DN/+^*) hearts at P14. (**D**) TRα binding site to the genomic locus of *Coq10a*, based on the TRα chromatin immunoprecipitation-sequencing (ChIP-seq) data. Values are reported as Mean ± SEM (n=3). *p<0.05, ***p<0.001.

To test if thyroid hormone signaling is necessary for *Coq10a* expression in the heart, we analyzed its transcript levels in mice that express a dominant negative form of thyroid hormone receptor alpha (Thra^DN^) to suppress thyroid hormone signaling specifically in cardiomyocytes (*Myh6-Cre;Thra^DN^*), recently generated by our laboratory (*19*). RNA-seq analyses of *Myh6-Cre;Thra^DN^* hearts at P14 demonstrated that the gene expression of *Coq10a* was reduced to one-third compared to control hearts (Figure 2B). Consistently, Coq10a protein was also reduced in *Myh6-Cre;Thra^DN^* hearts at P14, compared to the littermate control. Furthermore, our chromatin immunoprecipitation-sequencing (ChIP-seq) analysis revealed that thyroid hormone receptor alpha binds to the first intron of *Coq10a* (Figure 2D), thus both RNA-seq and ChIP-seq results provide strong supports for *Coq10a* as a direct target gene of thyroid hormone receptors. In sum, our findings demonstrate that thyroid hormone signaling activation is important for the cardiac gene expression of *Coq10a*.

### Mice lacking *Coq10a* develops cardiac hypertrophy at postnatal day 14

In fission yeast Coq10p, Phe39-Lys45 have been mapped as key CoQ-binding residues (*7*). In this sequence, Phe39 and Pro41 are two highly conserved amino acids from bacteria to human (Figure 3A) and have been shown by site-directed mutagenesis to be essential for CoQ interaction (*7*). To generate *Coq10a* knockout mice, we exploited the CRISPR/Cas9-based genome-editing technique and designed sgRNA targeting the upstream of the conserved ubiquinone-binding site (Figure 3B). Sequencing of one stable line of mutant mice at the F2 generation revealed an insertion of two nucleic acids at the targeted site, resulting in frameshift mutations, which produces a truncated Coq10a protein lacking the ubiquinone-binding site (Figure 3B). Coq10a protein expression was decreased in the *Coq10a^+/-^* heterozygous heart and was undetectable in heart of adult *Coq10a^-/-^* homozygous mutant mice (Figure 3C).

**Figure 3.**
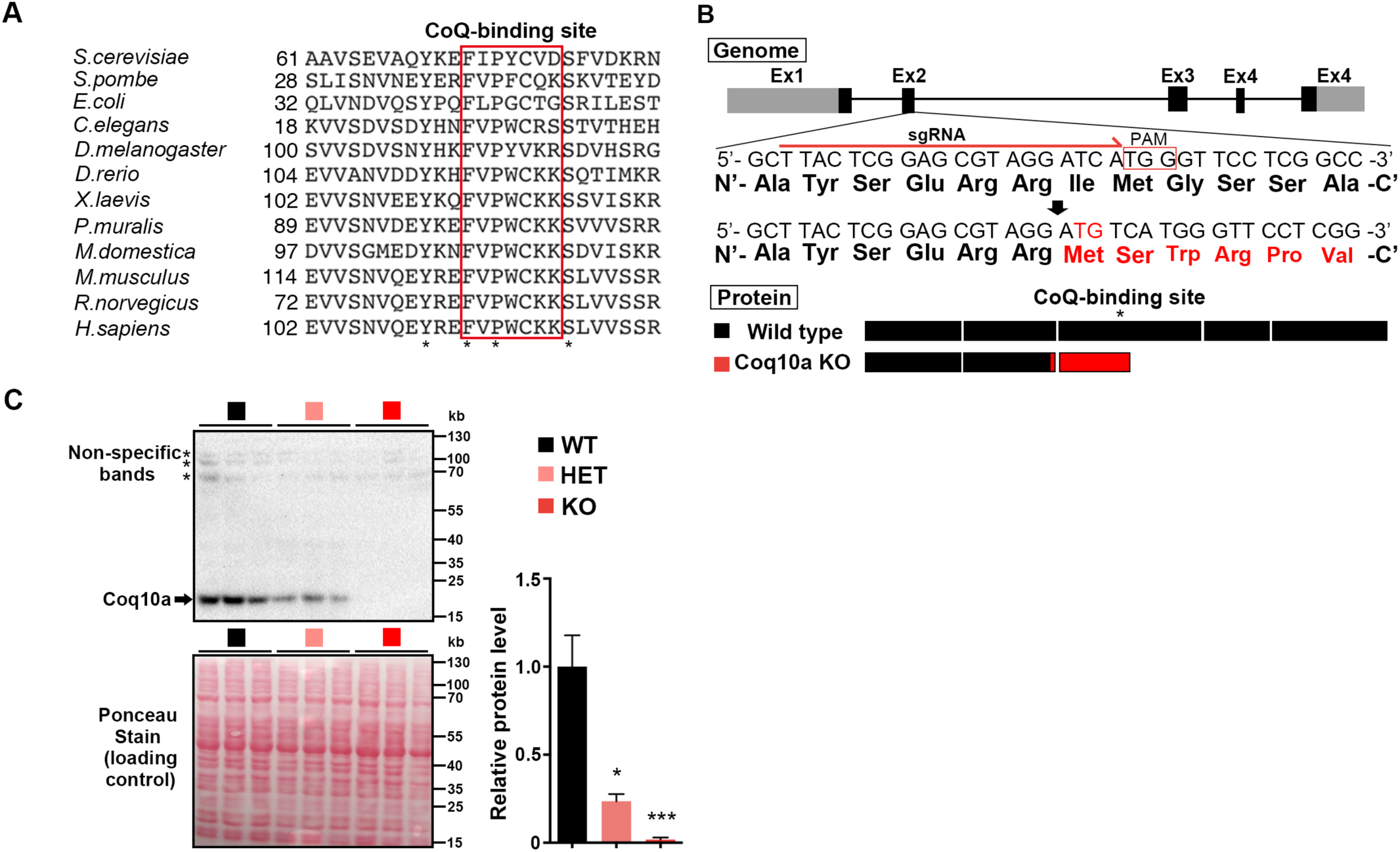
Generation of *Coq10a* knockout mice using CRISPR/Cas9. (**A**) Alignment of bacteria, yeast and vertebrate Coq10a protein sequences near the Coenzyme Q (CoQ)-binding site. The red box marked the conserved CoQ-binding site. (**B**) Schematic for generating *Coq10a* knockout mice. The sgRNA was designed to target a sequence on the second exon, and the mutant mice had an insertion of two nucleic acids, resulting in frameshift mutations. *Coq10a* knockout (KO) mice produce the truncated Coq10a protein lacking CoQ-binding site. (**C**) Western blotting analysis of Coq10a protein in adult wildtype (WT), heterozygous (HET) and KO hearts (n=3 animals). Values are reported as Mean ± SEM (n=3). *p<0.05, ***p<0.001.

*Coq10a^-/-^* mutant mice are born at Mendelian ratio with normal gross morphology and fertility. We first evaluated heart phenotypes at P14. *Coq10a^-/-^*mutant mice had an increase of the ratio of heart weight-to-body weight by 22% despite normal body weight (Figure 4A and 4B), indicating that depletion of *Coq10a* gene results in a cardiomegaly phenotype. This phenotype is further confirmed in seven F0 *Coq10a* homozygous mutant mice at P14 (Figure S2). Although gene editing using the CRISPR/Cas9 system might cause off-target effects, the highly reproducible cardiomegaly phenotype in F2 and multiple F0 mutant animals suggest that the phenotype is likely a consequence of the specific loss of *Coq10a*.

**Figure 4.**
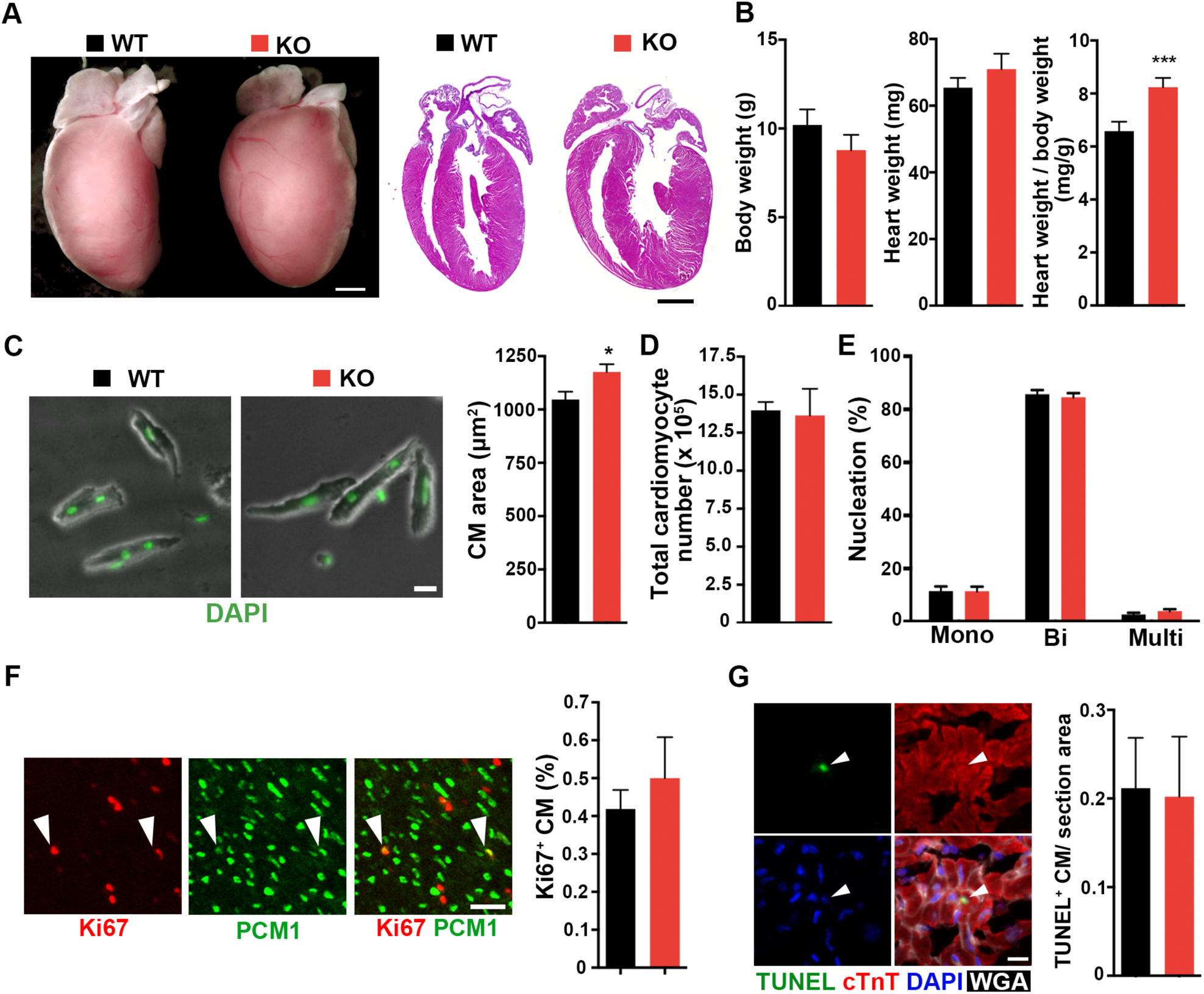
Cardiac hypertrophic phenotype of *Coq10a* knockout mice at P14. (**A**) Representative images of *Coq10a^-/-^* knockout (KO) and control wildtype (WT) hearts at P14. Whole-mount (left) and cross-section (right, hematoxylin-eosin stained) views. (**B**) Measurement of the bodyweight, heart weight and heart weight / body weight (n=9 animals). (**C**) Analysis of cell area of isolated cardiomyocytes (CM) from P14 heart (n=5 animals). (**D**) Total CM number in P14 hearts (n=4 animals). (**E**) CM nucleation in P14 heart (n=3 animals). Mono, Bi and Multi represent mononucleated, binucleated and multinucleated CMs, respectively. (**F**) Analysis of CM proliferative activity. Representative images of proliferating CMs (left). Arrowheads indicate Ki67^+^ CMs. Quantification of proliferating CMs that are double positive of Ki67 and PCM1 (n=3 animals) (right). (**G**) Analysis of apoptotic cells using TUNEL staining. Representative images of apoptotic CMs (left). TUNEL staining detects apoptotic cell (green). cTnT is a marker for CM cytoskeleton (Red). Nuclei were visualized by DAPI. Wheat Germ Agglutinin (WGA) labels cell boundary (white). Arrowhead indicates TUNEL+ CM (right). Quantification of TUNEL-positive CMs per section area (μm) (left) (n=3 animals). Values are reported as Mean ± SEM. *p<0.05, **p<0.01, Scale bars, 1mm (**A**), 20 μm (**C**), 50 μm (**F**) and 10μm (**G**).

To determine if the increased heart size is due to cardiomyocyte hypertrophy or hyperplasia, we first assessed the size of individual cardiomyocytes isolated from ventricle of P14 *Coq10a^-/-^*mice. Cardiomyocyte size was significantly increased in *Coq10a^-/-^*mice (Figure 4C). Next, to evaluate a possible contribution of hyperplasia to the cardiomegaly phenotype, we evaluated cardiomyocyte number, nucleation, proliferation and apoptosis in *Coq10a^-/-^*and control littermate mice at P14. Analyses of mutant and control mice showed comparable total cardiomyocyte numbers in the ventricle (Figure 4D) and cardiomyocyte nucleation distribution (Figure 4E). Proliferating cardiomyocytes was assessed by immunostaining against cardiomyocyte nuclear marker pericentriolar material 1 (PCM1) and a proliferation marker Ki67. However, we observed no difference in proliferating cardiomyocytes between *Coq10a^-/-^* mice and littermate controls (Figure 4F). To assess the extent of cell apoptosis, terminal deoxynucleotidyl transferase mediated deoxyuridine triphosphate nick end labeling (TUNEL) assays were performed on histological sections and no difference in the number of TUNEL-positive cardiomyocytes was observed between *Coq10a^-/-^* and control mice (Figure 4G). Therefore, our investigation suggests that the cardiomegaly is primarily due to increase in cardiomyocyte size, which is indicative of cardiac hypertrophy.

To understand how deletion of *Coq10a* led to heart hypertrophy, we performed RNA-seq analyses of *Coq10a^-/-^* and control hearts at P14. By gene ontology (GO) analysis, we found upregulated genes primarily in the categories of mitochondrion and metabolic process while downregulated genes mainly in the membrane, cell-cell junction and focal adhesion categories (Figure 5A). Moreover, gene-set enrichment analyses (GSEA) revealed citrate (TCA) cycle, Peroxisome Proliferator-Activated Receptor (PPAR) signaling pathway, fatty acid metabolism, and valine-leucine-isoleucine degradation pathway genes were significantly up-regulated (Figure 5B). In the genes down-regulated in *Coq10a^-/-^* mutant heart, we found an enrichment of components involved in extracellular matrix receptor interaction, dilated cardiomyopathy, DNA replication and hypertrophic cardiomyopathy (Figure 5B). Consistent with the upregulation of mitochondrial and metabolic genes, we found that *Ppargc1a* (also known as *Pgc1α*), *Pparg* and *Rxrg* expression significantly increased in the mutant heart (Figure 5C). These genes encode transcription factors that function as master regulators of mitochondrial biogenesis and energy metabolism.

**Figure 5.**
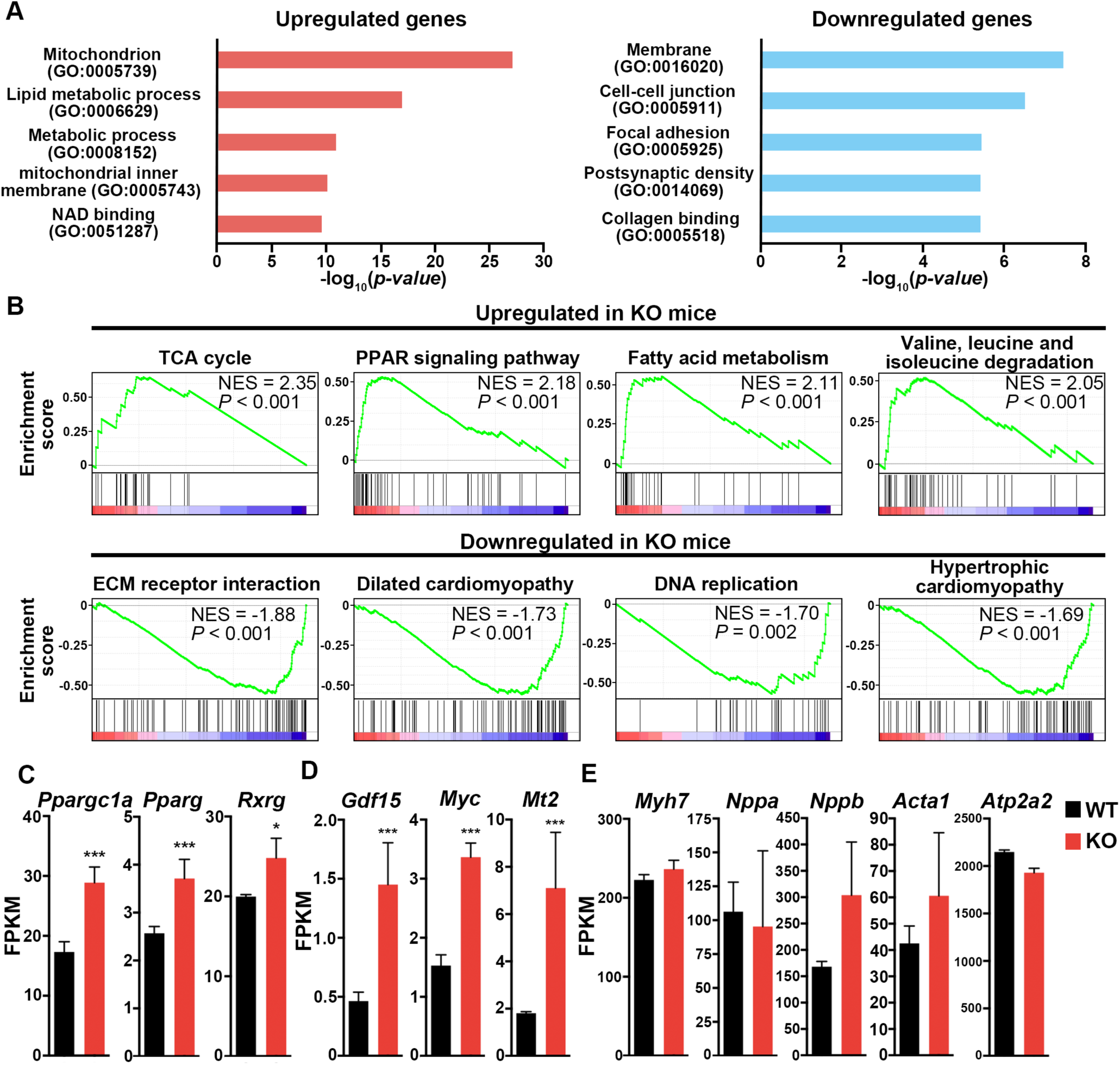
Transcriptomic analysis of *Coq10a* knockout heart at P14. (**A**) Gene ontology analysis of upregulated and downregulated gene categories in the mutant heart at P14 based on RNA-seq analysis. (**B**) Gene set enrichment analysis of upregulated and downregulated pathways in the mutant heart at P14. (**C**) Expression of mitochondria biogenesis-related genes. (**D**) Expression of cardiac hypertrophy and related genes. (**E**) No expressional change of typical pathological hypertrophy-induced fetal genes. Values are reported as Mean ± SEM (n=3 hearts). *p<0.05, ***p<0.001.

Among the genes whose expressions are significantly altered in the *Coq10a^-/-^* mutant heart, we identified *Growth/differentiation factor 15* (*Gdf15*), *c-Myc (myc)*, *Metallothionein 2* (*Mt2*) (Figure 5D) that have been previously shown to either induce cardiomyocyte hypertrophic growth or protect the heart against oxidative stress in hypertrophy (*20–23*). It was demonstrated that GDF15 proteins increased the rate protein synthesis by 47% and cardiomyocyte hypertrophic growth by 27% *in vitro* (*20*). Mice with inducible expression of *c-Myc* in post-mitotic cardiomyocytes develop cardiac hypertrophy through enhanced free fatty acid utilization and oxidative metabolism (*21*). *Mt2* is highly up-regulated in pregnancy-induced cardiac hypertrophy and overexpression of *Mt2* in mice has protective effects on cardiac oxidative stress through the antioxidant property of Metallothioneins (*22, 23*).

We next examined our RNA-seq data for possible gene expression signature of pathological cardiac hypertrophy. Development of pathological hypertrophy is accompanied by up-regulation of fetal heart genes such as *Acta1*, *Myh7*, *Nppa*, *Nppb* and *Atp2a2*. The expression of these genes, however, is not significantly activated in our mutant mice (Figure 5E). In pathological hypertrophic heart, cardiac fibroblasts differentiate into myofibroblasts that produce type I collagen to progress cardiac fibrosis. The expression level of alpha-smooth muscle actin (α-SMA) (*Acta2*), which is known as a marker for differentiation of fibroblasts into myofibroblasts was not increased in our mutant mice. Consistently, no increase of type I collagen gene expression (*Col1a1* and *Col1a2*) was observed in *Coq10a^-/-^* heart (Figure S3B). Moreover, pathological hypertrophy is commonly accompanied by myocardial cell death. We did not observe any increased expression of executioner caspase genes (*Casp3, 6* and *7*) or apoptosis associated genes (*Bax* and *Bcl2*) in our mutant heart (Figure S3A), which is consistent with the TUNEL analysis (Figure 4G). In addition, the expression of Coq family genes including *Coq10b* was not altered in *Coq10a* mutant (Figure S4), indicating that there was no compensational upregulation of other Coq family genes.

### Adult *Coq10a* knockout mice maintain cardiac hypertrophy with preserved cardiac functions and mitochondrial respiration

Next we analyzed cardiac phenotypes of *Coq10a* knockout mice in adulthood. Compared to littermate wild type, the ratio of heart weight to body weight in adult *Coq10a^-/-^* heart increased 25 % (Figure 6A), which is comparable with the extent of heart size change at P14. Consistent with comparable expression of collagen type I genes at P14, adult *Coq10a^-/-^* mutant hearts did not display significant fibrosis (Figure 6B). Furthermore, echocardiography showed similar contractile functions between *Coq10a* knockout and control heart, suggesting preserved cardiac function in *Coq10a^-/-^*mice (Figure 6C). Even in eight months old mutant mice, we did not observe any defects in cardiac function. In addition, blood pressure and heart rate measurements indicate that *Coq10a^-/-^*heart functions normally in terms of systolic blood pressures and heart rate (Figure 6D). Taken together, our results suggest that knockout of *Coq10a* leads to transcriptional up-regulation of hypertrophy-inducing genes without any features of pathological hypertrophy or heart failure.

**Figure 6.**
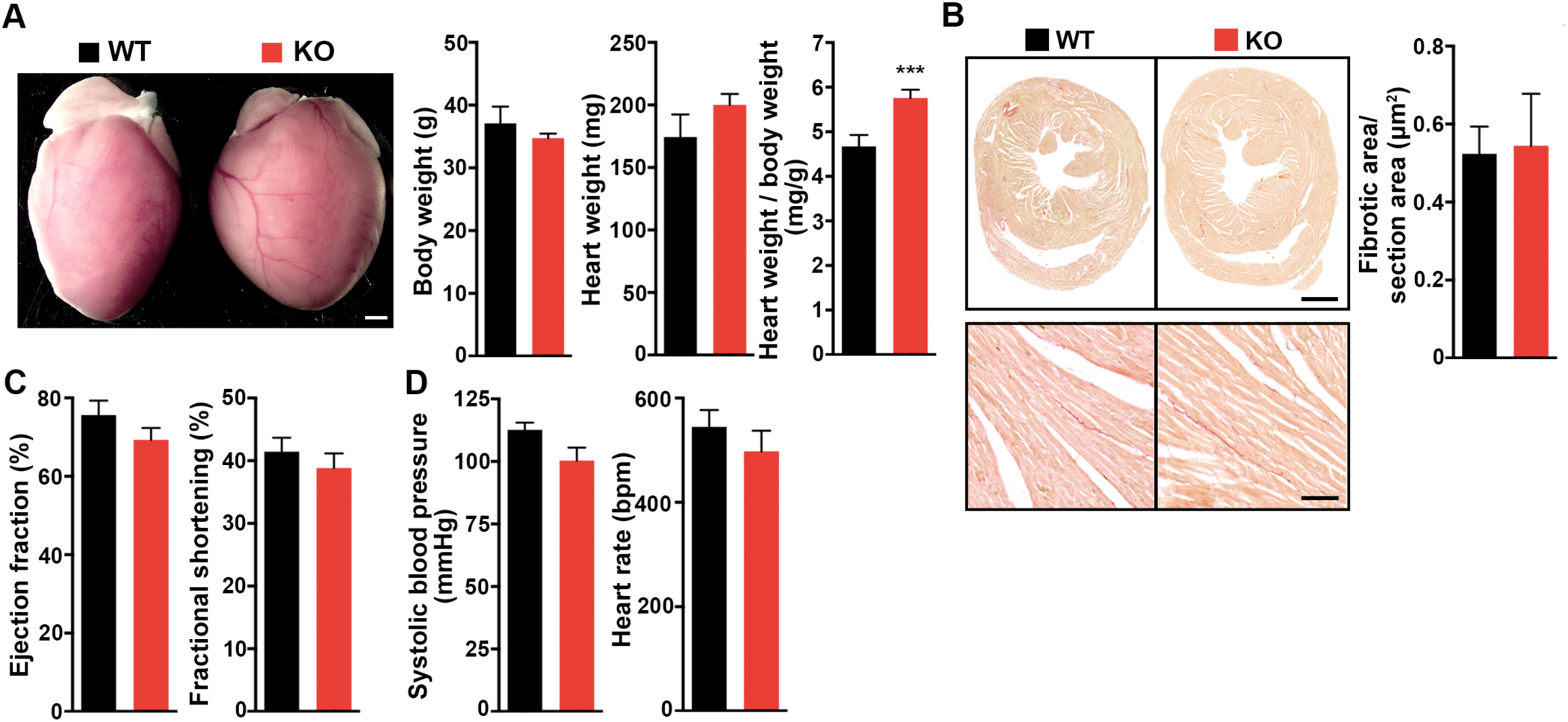
Cardiac hypertrophic phenotype of *Coq10a* knockout mice in adulthood. (**A**) Representative whole-mount images of adult *Coq10a* knockout (KO) and littermate wildtype (WT) hearts (left). Measurement of body weight and heart weight (right) (n=9 animals). (**B**) Fibrosis analysis. Sirius red staining of heart sections (left). Quantification of fibrotic area per section area (μm^2^) (right). Animals are 5-8 month old. (**C**) Measurement of cardiac functions by means of echocardiography. The ejection faction and fraction shortening data of individual animals (5-8 month old) are presented (n=6-8 animals). Measurements of systolic blood pressure and heart rate. (**D**) No difference of blood pressure and heart rate was observed between WT and KO animals (n=6-8 animals). Values are reported as Mean ± SEM. ***p<0.001, Scale bars, 1mm (**A**), 1mm in upper panel and 100 μm in lower panel (**B**).

Yeast Coq10p has been hypothesized to be a CoQ binding protein which transports CoQ from the synthase complex to proper positions in the mitochondrial inner membrane (*4*). Mutations of *COQ10* in yeast cause decreased or normal level of coenzyme Q, with mitochondrial respiration defects (*4*). To investigate the role of *Coq10a* in mitochondrial biogenesis and function, we first isolated mitochondria from *Coq10a^-/-^* mutant and control hearts. Intriguingly, the total amount of protein from purified mitochondria per ventricle significantly increased from 565 ± 52.9 mg in control hearts to 959 ± 58.4 mg in mutant hearts (Figure 7A). Next, we measured the oxygen consumption rate (OCR) in isolated mitochondria but found no significant difference between *Coq10a^-/-^* and littermate control mice (Figure 7B). This result suggests that in contrast to yeast *coq10* mutants, *Coq10a^-/-^* mutant mice have intact mitochondrial respiration in the heart (Figure 7B). Furthermore, we analyzed the levels of CoQ_9_ (the main coenzyme Q specie in mice) and CoQ_10_ by high performance liquid chromatography (HPLC) and demonstrated 1.7-fold increase of CoQ_9_ and 2.1-fold increase of CoQ_10_ in *Coq10a^-/-^* hearts, compared to control (Figure 7C). Consistently, we found that the ratio of mitochondria DNA to nuclear DNA were almost doubled in the *Coq10a^-/-^* heart (Figure 7D), suggesting that deletion of *Coq10a* may result in an accumulation of functional mitochondria and coenzyme Q in the mouse heart. The increase of mitochondrial proteins and DNA could be explained by upregulation of *Pgc1α* gene expression in the *Coq10a^-/-^* heart (Figure 5C). PGC1α is a master regulator of mitochondrial biogenesis and energy metabolism. Cardiac-specific overexpression of *Pgc1α* to a non-physiological high-level leads to pathological cardiac hypertrophy followed by uncontrolled mitochondrial biogenesis and loss of sarcomere structure (*24, 25*). Although the expression of *Pgc1α* and its downstream genes are generally downregulated in pathological hypertrophy (*26*), exercise-induced physiological hypertrophy shows moderate increases (approximately 50-80%) of *Pgc1α* expression (*27–31*). Thus, the 67% increase of *Pgc1α* mRNA in the *Coq10a^-/-^* heart (Figure 5C) would be consistent with the mutant heart and mitochondrial phenotypes. In sum, our findings suggest that depletion of *Coq10a* increases mitochondrial content with preserved mitochondrial respiration in the heart possibly through PGC1α pathway activation.

**Figure 7.**
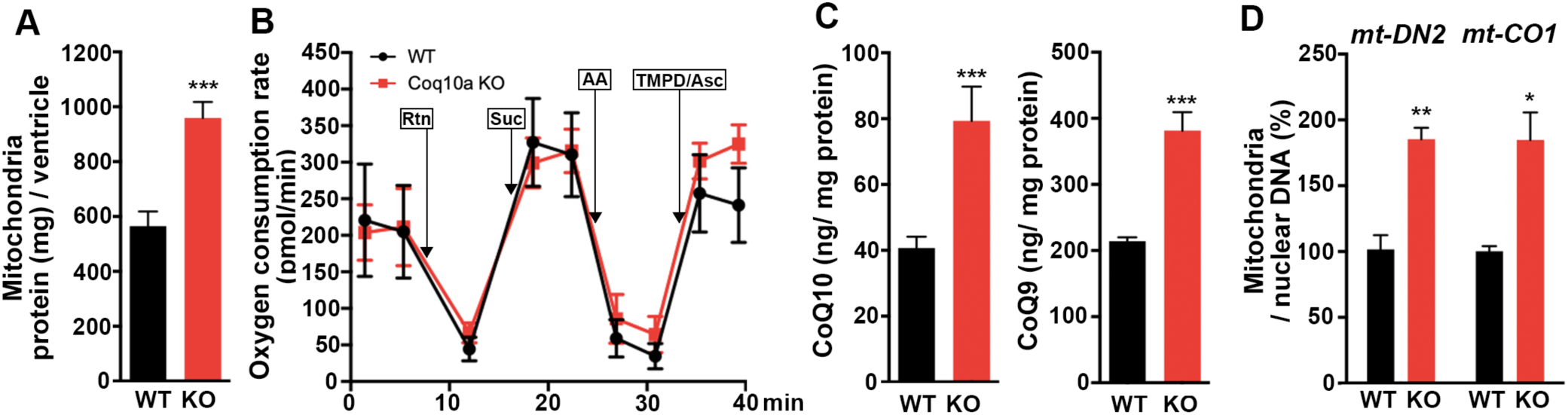
Mitochondrial activity in *Coq10a* knockout mice. (**A**) Quantification of mitochondrial protein per ventricle in adult mice. (**B**) Quantification of oxygen consumption rate in isolated mitochondria (n = 4 to 5 animals). Arrows indicate the time of drug injections. Rtn, rotenone. Suc, succinate. AA, antimycin A. TMPD/Asc, TMPD/ascorbate. (**C**) Measurements of CoQ_10_ and CoQ_9_ per total ventricular protein (mg) (n = 4 to 5 animals). (**D**) Mitochondrial DNA analysis (n=4 hearts). Values are reported as Mean±SEM. *p<0.05, **p<0.01, ***p<0.001.

## Discussion

In this report, we used a genetic approach to investigate the physiological function of the novel mitochondrial protein Coq10a. Mice lacking *Coq10a* develop cardiac hypertrophy without showing any pathological hypertrophic phenotypes at P14 and in adulthood. Despite mice lacking various mitochondria-related gene often develop pathological hypertrophy due to dysfunctional mitochondria (*11*), *Coq10a* knockout mice have normal mitochondrial respiration and increased mitochondrial content, consistent with increased expression level of *Pgc1α* which encodes a master regulator of mitochondrial biogenesis. In addition, *Coq10a* knockout mice have increased levels of CoQ in the heart.

Based on our analyses, the hypertrophic phenotype of *Coq10a*-deleted heart seems to be different from pathological hypertrophy which occurs as a consequent of deleterious genetic mutations or in response to hypertension and ischemic injury (*32, 33*). Pathological hypertrophy is often accompanied by progressive contractile dysfunction and eventually heart failure. The decrease of mitochondrial gene expression followed by reduced mitochondrial respiration often develop in pathological hypertrophy (*33*). In contrast, physiological hypertrophy is a beneficial phenotype, which develop during postnatal development, pregnancy and exercise (*32*). Unlike pathological hypertrophy, it increases or preserves cardiac functions with enhanced mitochondrial biogenesis (*33*). Thus, our reported preserved cardiac contractile and mitochondrial functions in the *Coq10a* mutant mice suggest that the observed hypertrophic phenotype is more likely an enhanced physiological hypertrophy during the postnatal growth.

The mitochondrial phenotypes of *Coq10a* knockout mice generated in this study are different from those of *coq10* yeast mutants. In yeast, the CoQ biosynthetic protein complex assembles at ER-mitochondria contact sites (*1, 2*), and yeast Coq10p protein is hypothesized to be a carrier of CoQ to its proper location in the inner mitochondrial membrane for functioning in the electron transport chain (*4*). Accordingly, yeast lacking *COQ10* gene present defects in mitochondria respiration (*4, 5*). In contrast, the mitochondria of *Coq10a* knockout heart showed normal respiration activity when normalized by the total mitochondrial protein level. Given that *Coq10a* mutant mice have more mitochondrial proteins but normal cardiomyocyte numbers per heart, we predict that mice lacking *Coq10a* would have enhanced mitochondria respiration activity per cardiomyocyte. Additionally, despite yeast *coq10* mutant shows normal levels of CoQ, *Coq10a* mutant mice have 2-fold increases of CoQ levels. RNA-seq analyses of P14 hearts indicates no compensatory up-regulation of *Coq10b* and genes involved in CoQ synthesis (Figure S4), suggesting that the increase of CoQ levels might be independent of enhanced CoQ synthesis. One possible explanation for the increase of CoQ levels is that mouse Coq10a protein may work as a carrier of CoQ as what was suggested for yeast Coq10p (*4*), and CoQ is accumulated in the mitochondria of *Coq10a* mutant mice because a significant proportion of CoQ is not transported and used properly. It is still an enigma why *Coq10a* mutant mouse has a normal respiration rate in mitochondria unlike yeast *coq10* mutant. The more ubiquitously expressed Coq10 member *Coq10b* (Figure S1) may play a redundant function with *Coq10a* to fulfill the role of yeast *COQ10* in mammalian mitochondria. Further investigation of the physiological function of *Coq10b in vivo* will help test this possibility.

The molecular mechanism by which deletion of the mitochondrial protein Coq10a affects transcription of genes such as *Pgc1α* is still unknown. However, there are some evidences showing supplementation of CoQ_10_ increases *Pgc1α* expression and activates PPAR pathway in mouse liver (*34, 35*). In addition, coenzyme Q10 has recently been reported to be a novel potent ligand for PPARα/γ (*36*). Since our mutant mice have a 2.1-fold of CoQ_10_, there is a possibility that the accumulation of CoQ triggers mitochondrial biogenesis via direct activation of PPARα/γ in our mutant mice. Nevertheless, understanding of how Coq10a regulates cardiomyocyte mitochondrial gene expression could yield new insights into the molecular control of coenzyme Q post-synthesis transport and postnatal heart hypertrophic growth.

## Methods

### Animals

Mice procedures were conducted in accordance with the Institutional Animal Care and Use Committee (IACUC) of the University of California, San Francisco. CD-1 (Charles River) and *Coq10a*^-/-^ mouse lines were maintained according to the University of California, San Francisco institutional guidelines. Adult mice were 2-8 month old. For all other experiments, both male and female animals were used and no gender difference was observed. Generation of *Myh6-Cre;Thra^DN/DN^* mice, RNA-seq analysis in *Myh6-Cre;Thra^DN/DN^* mice and ChIP-seq analysis against *Thra-GFP* were described in our previous paper (*19*).

### Western blotting

Proteins were lysed in 1% SDS (reconstituted in PBS) supplemented proteinase inhibitor (Fisher, PI88666) from frozen P14 mouse heart ventricles. Protein concentration was measured using the BCA Protein Assay Kit (Pierce). 100mg protein lysate was first loaded to a NuPAGE 4-12% Bis-Tris Gel (Invitrogen) for separation, and then transferred to a PVDF membrane (BIO-RAD). Protein loading was assessed using Ponceau S staining. The blot is visualized using ECL Western Blotting substrate (Pierce). The following primary antibodies were used: rabbit anti-Coq10a (Proteintech, 17812-1-AP, 1:1000), mouse anti-Troponin T (Thermo Fisher Scientific, MS295P1, 1:1000), mouse anti-Tom20 (Santa Cruz Biotechnology, sc-17764, 1:1000).

### Gene expression analysis on neonatal CMs

For CM isolation, mice were anesthetized on ice, and then the freshly isolated heart was cannulated onto a needle and perfused with 2 mg/mL collagenase II (Worthington) until the heart loses integrity. The isolated CMs were plated on 96 well plate coated by Laminin in 5% FBS in DMEM supplemented with 1μM Ara-C and 100 μg/ml primocin. At 24 hours after the culture, the media was replaced to 0.05% BSA in DMEM supplemented with 100 μg/ml primocin, and 30nM or 300nM T3 were added. At 72 hours after culture, total RNA was isolated using TRIzol, and then cDNA was synthesized using iScript™ cDNA Synthesis Kit. qPCR was performed using the SYBR Select Master Mix (Applied Biosystems, 4472908) and the 7900HT Fast Real-Time PCR system (Applied Biosystems). Relative mRNA levels of *Coq10a, Myh6 and Actb1* were measured using the following primers: *Coq10a* Forward = AGCGAAAGGCTTACTCGGAG; *Coq10a* Reverse = TGGACACCACCTCAAACATCT; *Myh6* Forward = TGCACTACGGAAACATGAAGTT; *Myh6* Reverse = CGATGGAATAGTACACTTGCTGT; *Actb1* Forward = AGTGTGACGTTGACATCCGT; *Actb1* Reverse = TGCTAGGAGCCAGAGCAGTA.

### Sequence alignment

Protein sequences were obtained from NCBI. The following shows the protein accession numbers of the sequences in variant species. These sequences were selected based on similarity to mouse *Coq10a* using BLAST. *S.cerevisiae* (NP_014635.1), *E.coli* (WP_137445723.1), *S.pombe* (NP_587994.1), *C.elegans* (NP_001040866.1), *D.melanogaster* (NP_724484.1), *D. rerio* (XP_009295305.1), *X.laevis* (XP_018105495.1), *P.muralis* (XP_028565875.1), *M.domestica* (XP_001379337.1), *M.musculus* (NP_001074509.1), Rattus norvegicus (NP_001102197.1), and *H.sapiens* (NP_653177.3). The sequence alignment was generated using Clustal Omega.

### Generation of Coq10a knockout mice

The sequence of sgRNA targeting *Coq10a* is TTACTCGGAGCGTAGGATCA. CD-1 (Charles River) was used to generate knockout mice. Purified sgRNA and Cas9 protein were injected into mouse-fertilized oocytes by CVRI Mouse Core Facility.

### Histology

3.7% paraformaldehyde (PFA) fixed hearts were embedded in paraffin, and then sectioned 5 μm thickness. For fibrosis staining, sections were soaked in Sirius Red for 25 min at room temperature. For eosin and hematoxylin staining, sections were stained with Eosin for 3 min and hematoxylin for 5 min at room temperature. Sections were imaged using Axio Scan.Z1 and fibrotic area was quantified using ImageJ.

### Cardiomyocyte isolation

Mice were injected with 6.67 mg/kg bodyweight of heparin (1000 IU/mL) intraperitoneally 30 minutes before anesthetizing the mice with 20% ethyl-carbamate. The hearts were rapidly excised and the aorta was cannulated onto a Langendorff apparatus and perfused with 2 mg/mL collagenase II (Worthington). Digestion was stopped after the hearts lost partial integrity. Hearts were then unmounted from the Langendorff apparatus, ventricles isolated, and gently triturated with forceps to further release cardiomyocytes. Isolated cardiomyocytes were then fixed in 2% PFA for 15 minutes at room temperature, spotted onto glass slides, and stained with DAPI to visualize nuclei. Cardiomyocyte size and nucleation were analyzed using ImageJ. At least 200 cells were used for the quantifications.

### Immunohistochemical staining on mouse heart

Methods for sectioning and immunostaining were described in our previous paper (). Briefly, freshly dissected heart was embedded in O.C.T. Compound and flash frozen on metal box in liquid nitrogen. Embedded samples were then sectioned with a Leica CM3050S to 5 μm thickness. PFA-fixed sections were permeabilized in 0.2% Triton X-100 in PBS, blocked in 5% normal donkey serum (NDS) in PBST, and incubated with primary antibodies. The following primary antibodies were used: rat anti-Ki67 eFluor 570 (eBiosciences, 41-5698-80, 1:200), rabbit anti-PCM1 antibody (Sigma, HPA023370, 1:2000) and mouse anti-Troponin T (Thermo Fisher Scientific, MS295P1, 1:200) in PBST overnight at 4°C. After washing with PBST, samples were incubated in secondary antibodies with fluorescence (1:500) for 2 hours at room temperature. Cell membrane was labeled with wheat germ agglutinin (1:200) and nuclei were stained with DAPI. TUNEL staining was followed the instruction accordingly.

### Quantification of total cardiomyocyte number

Hearts were fixed in 3.7% formaldehyde for 1 hour. After removal of the atria, the ventricle was then cut into four pieces and digested in PBS containing 4 mg/ml collagenase II (Worthington) with gentle shaking at 37°C until complete digestion. The digestion buffer was changed daily. All dissociated cardiomyocytes were pooled and counted using a hemocytometer.

### RNA-seq analysis on Coq10a KO mice

RNA was isolated from heart samples using TRIzol Reagent (Thermo Fisher Scientific) according to a manufacturer’s instruction and further pre-cleaned by Zymo RNA Clean & Concentrator 5. 1 mg of total RNA was used for library preparation with Illumina Tru-seq RNA Library Prep Kit V2. NEXTflex Unique Dual Index Barcodes (PerkinElmer) were used to minimize index hopping. Samples were normalized, pooled and sequenced on Illumina NovaSeq 6000 to obtain 50 bp paired-end reads.

### Measurements of cardiac functions

Left ventricular systolic function was evaluated by two-dimensional echocardiography with Visual Sonics Vevo 3100 equipped with a 40 MHz probe. Animals were anesthetized with 0.5-1.0% isofluorane and hair was removed over the measurement area. The mice were then placed in a supine position on a heating pad. To measure ejection fraction and fractional shortening, short axis images were acquired at the level of the papillary muscle with M-mode, and the left ventricle internal dimensions (LVIDs) were determined in both diastole and systole.

Blood pressure and heart rate were measured using BP-2000 Blood Pressure Analysis System^TM^ (Visitec Systems). The values were determined by average numbers of experimental data in three days.

### Quantification of mitochondrial DNA

DNA was purified from frozen P14 heart ventricles with Proteinase K, following by phenol/chloroform extraction. qPCR was performed using the SYBR Select Master Mix (Applied Biosystems, 4472908) and the 7900HT Fast Real-Time PCR system (Applied Biosystems). Mitochondrial DNA content was measured via qPCR targeting mitochondrial genes, *ND2* and *Cox1*, and Relative mitochondrial DNA content was measured with qPCR targeting mitochondrial genes, *ND2* and *Cox2*, and normalization against genomic gene, *H19* using the following primers: *mtND2* forward=5’- CCCATTCCACTTCTGATTACC-3’; *mtND2* reverse= 5’- ATGATAGTAGAGTTGAGTAGCG-3’; *mtCox1* forward =5’- CTGAGCGGGAATAGTGGGTA-3’; *mtCox1* reverse= 5’- TGGGGCTCCGATTATTAGTG-3’; *H19* forward =5’- GTCCACGAGACCAATGACTG-3’ ; *H19* reverse =5’-GTACCCACCTGTCGTCC-3’.

### Separation of mitochondrial and cytosol fractions

Mitochondria Isolation Kit for Tissue (Abcam, ab110168) was used for the separation of mitochondrial and cytosol fractions. Freshly isolated whole heart was removed atria, washed with pre-chilled washing buffer, and minced in pre-chilled isolation buffer. After spinning down at 1,000g, supernatant was transferred to new tube, and then was centrifuged at 12,000g for the separation. Supernatant containing cytosol protein was mixed with 1% SDS solution supplemented with proteinase inhibitor (PI) cocktail (Roche, 5892970001), and then was used for western blotting. The sedimented mitochondria were washed in isolation buffer supplemented with PI cocktail (Abcam, ab201111), and was used for mitochondrial respiration analysis. For western blotting, the mitochondria were dissolved in 1% SDS solution supplemented with PI cocktail (Roche, 5892970001). The protein concentration was determined using BCA assay.

### Measurement of mitochondrial respiration

Seahorse Bioscience XFe24 Extracellular Flux Analyzer was used for the measurement of mitochondria respiration. 5μg of mouse heart mitochondria was plated in each wells of Seahorse XF24 24-well microplate, and added 450μl of 1X MAS containing 4% BSA, 10 mM pyruvate and 5 mM malate. Isolated mitochondria were treated with 2 μM rotenone, 10 mM succinate, 5 μM antimycin A and 100 μM cN,N,N’,N’-Tetramethyl-p-phenylenediamine (TMPD) with 10mM ascorbate (Asc) at the indicated time points.

### Quantifications of CoQ_9_ and CoQ_10_

The detail method is described previously (*37*). CoQ_9_ and CoQ_10_ were extracted from ventricle in Coq10a KO mice and littermate control mice. The lipid component of the extract was separated using high-performance liquid chromatography (HPLC). CoQ concentration was measured by comparison of the peak area with those of standard solutions of known concentration and expressed in micrograms per gram of protein.

### Statistical analysis

The number of samples per each experimental condition is listed in the description of the corresponding figure legend. Statistical significance was determined using Student’s T-test for the figures. Error bars are represented as standard error of the mean.

## Acknowledgements

We thank Huang lab members for discussion. This work is supported by JSPS Overseas Research Fellowships (to K.H.), NIH (R01HL13845) Pathway to Independence Award (R00HL114738), Edward Mallinckrodt Jr. Foundation, March of Dimes Basil O’Conner Scholar Award, American Heart Association Beginning Grant-in-Aid, American Federation for Aging Research, Life Sciences Research Foundation, Program for Breakthrough Biomedical Research, UCSF Eli and Edythe Broad Center of Regeneration Medicine and Stem Cell Research Seed Grant, UCSF Academic Senate Committee on Research, REAC Award (Harris Fund), Department of Defense, and Cardiovascular Research Institute (to G.N.H.).

## Author contributions

K.H., S.C., H.Y., J.W., EB., X.C., S.K., K.T., T.Y. and G.H. performed experiments. K.H, X.C., S.K. analyzed data. S.K., C.M.Q. and G.H. provided reagents and contributed to discussions. K.H., and G.N.H. designed experiments and wrote the manuscript. The authors declare no competing interests.

---

**Figure S1.**
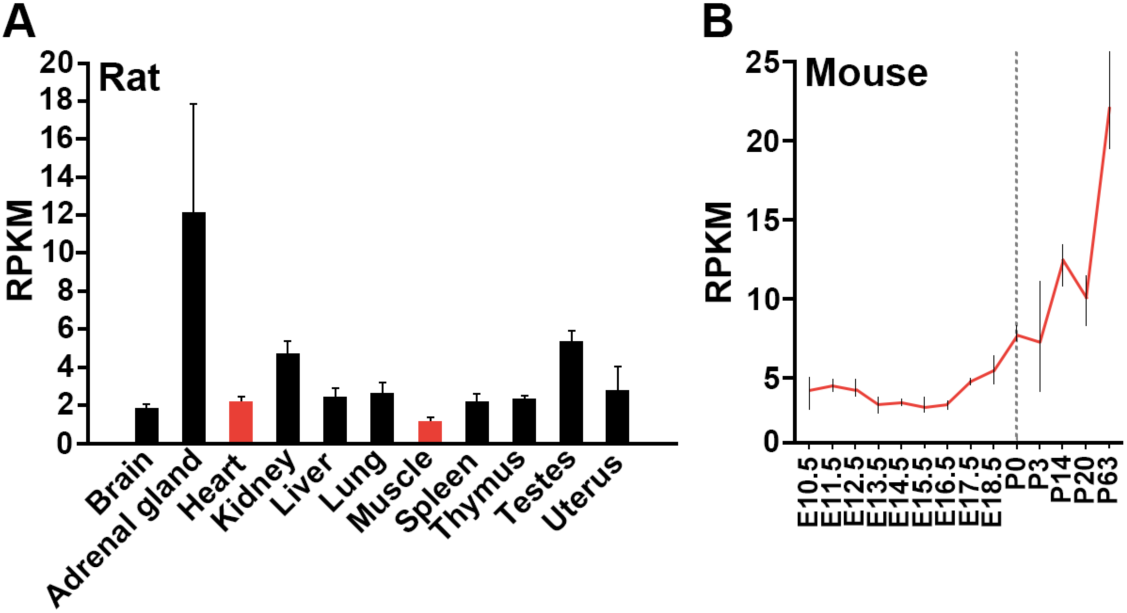
Tissue-specific and temporal expression of *Coq10b* gene in rodents. (**A**) *Coq10b* expression in eleven rat organs at 21 weeks old, based on the transcriptomic BodyMap (*14*). (**B**) Temporal *Coq10b* expressional changes from embryonic day (E) 10.5 to postnatal day (P) 63 in the mouse heart based on RNA-seq gene expression profile (*15*).

**Figure S2.**
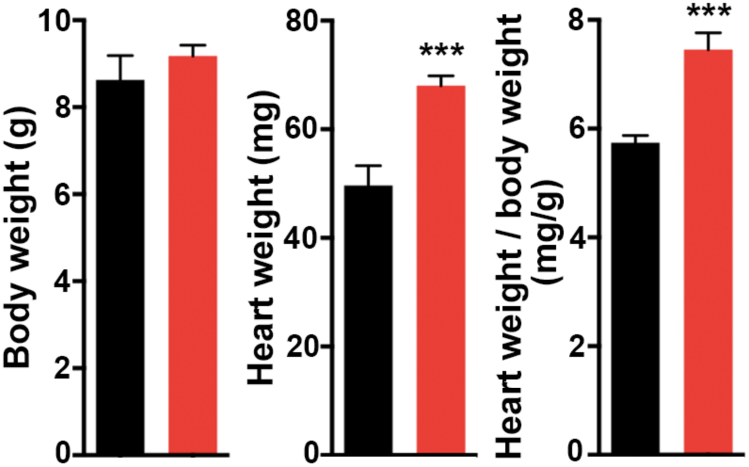
Cardiac hypertrophic phenotype of F0 *Coq10a* knockout mice at P14. Measurements of bodyweight, heart weight and heart weight / body weight (n=7-8 animals). Values are reported as Mean ± SEM. ***p<0.001.

**Figure S3.**
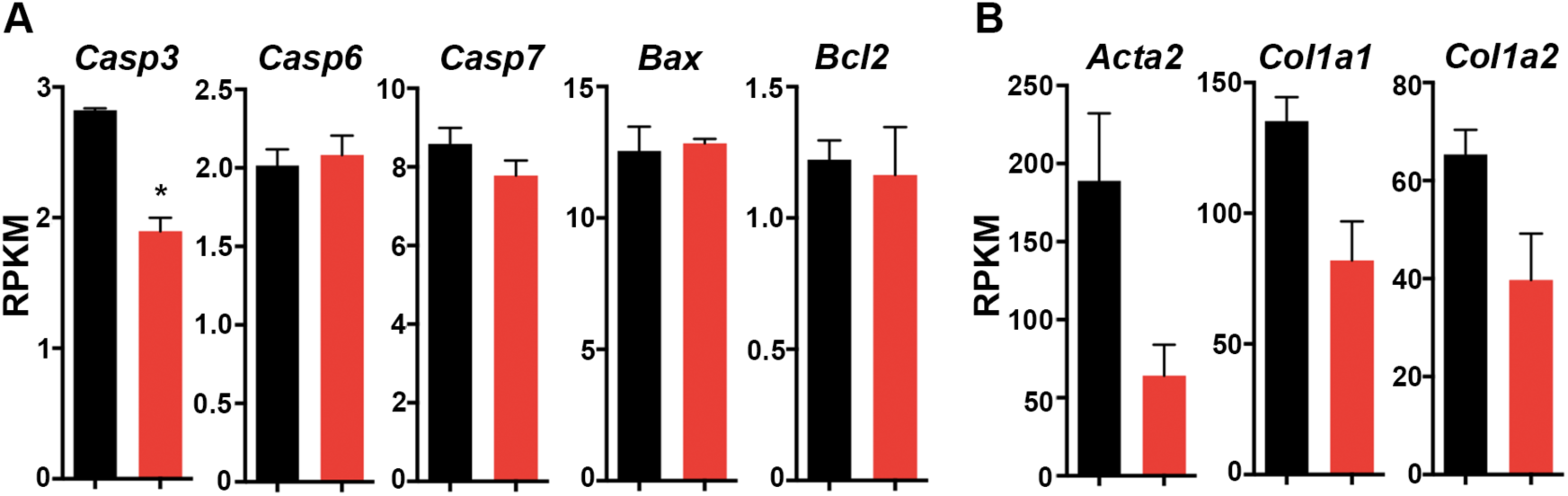
Gene expression of apoptosis and fibrosis-related genes in *Coq10a* knockout mice at P14. **(A)** Expression of apoptosis-related genes. **(B)** Expression of cardiac fibrosis-related genes, Values are reported as Mean ± SEM (n=3 hearts). *p<0.05.

**Figure S4.**
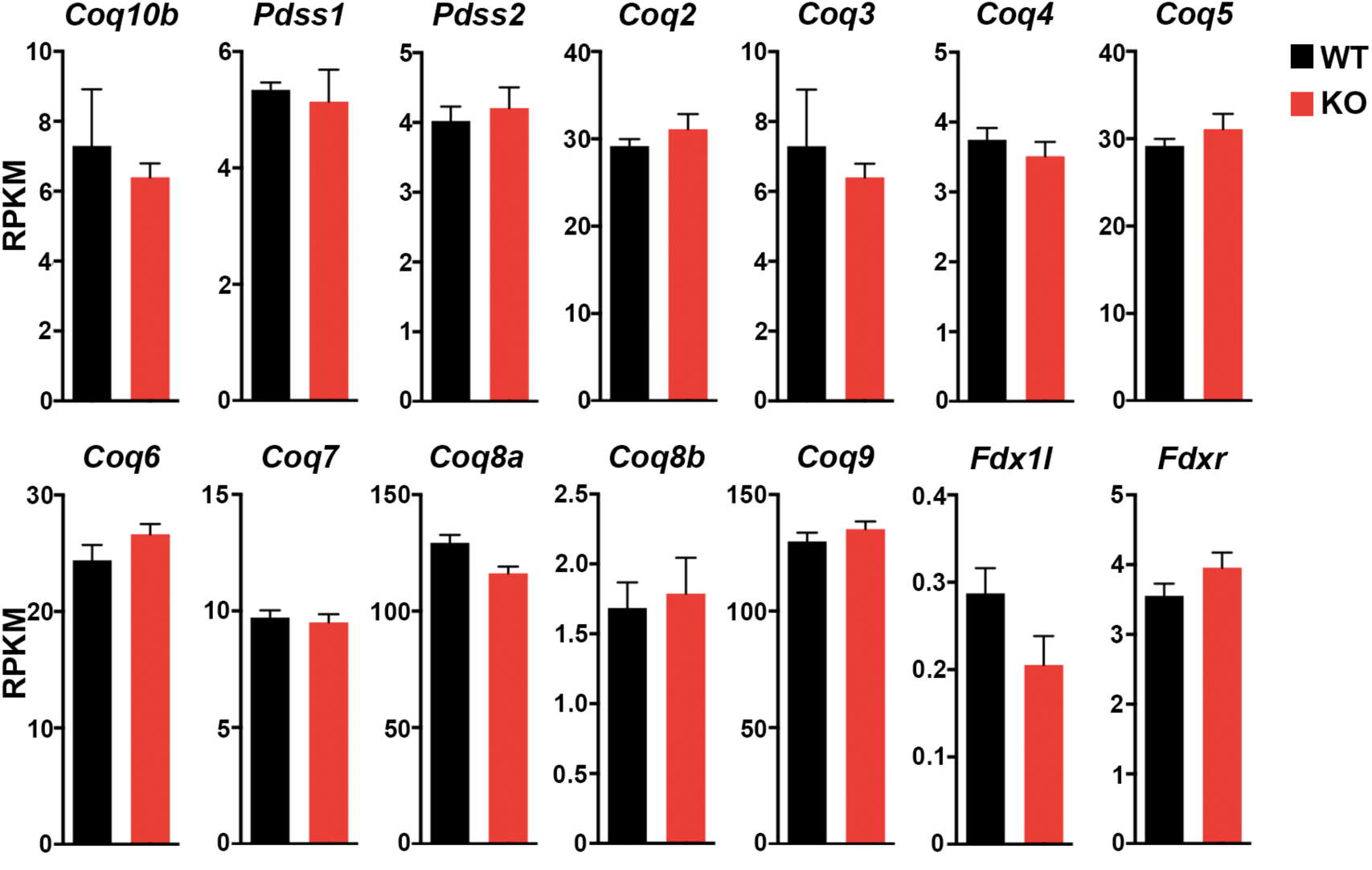
Expression of Coq family genes in *Coq10a* knockout mice at P14. Expression of Coq family genes. Values are reported as Mean ± SEM (n=3 animals).

